# Cefdinir binding to a class-A *β*-lactamase revealed by serial cryo-crystallography

**DOI:** 10.64898/2025.12.11.693660

**Authors:** Gargi Gore, Andreas Prester, Kim Bartels, David von Stetten, Eike C. Schulz

**Affiliations:** University Medical Centre Hamburg-Eppendorf (UKE), Hamburg, Germany; Institute for Nanostructure and Solid State Physics, University of Hamburg, Hamburg, Germany; European Molecular Biology Laboratory (EMBL), Hamburg, Germany; Max-Planck-Institute for the Structure and Dynamics of Matter, Hamburg, Germany

**Keywords:** serial crystallography, CTX-M-14, Cefdinir cephalosporins, antibiotic resistance

## Abstract

One of the most common resistance mechanisms against antibiotics employed by Gram-negative bacteria involves the production of *β*-lactamases, resulting in rapid hydrolysis of the antibiotic. Extensive use of the early generation cephalosporins led to the rise of extended-spectrum *β*-lactamases (ESBLs) like CTX-Ms. Cefdinir is an extended-spectrum third-generation cephalosporin administered since the late 90s; despite this, there is no reported 3D-structure of the antibiotic bound to any *β*-lactamase or Penicillin-Binding-Protein (PBP) in the PDB. Here we report the X-ray crystallographic structure of Cefdinir-bound CTX-M-14 E166A mutant obtained via serial cryo-crystallography (cryo-SSX).

**Synopsis:** Serial cryo-crystallography reveals the structure of the extended spectrum *β*-lactamase CTX-M-14, in complex with the third-generation cephalosporin antibiotic Cefdinir.

## 1 Introduction

Widespread antibiotic resistance is turning into a public health issue, claiming millions of lives annually (Ikuta *et al*., 2022; Murray *et al*., 2022). An alarming study showed that, in 2019 alone, almost 2 million deaths could be attributed to multidrug-resistant bacteria (Murray *et al*., 2022). Various pathogenic bacteria can employ a repertoire of different resistance mechanisms, including but not limited to: active expulsion of the antibiotic from the cell, generation of alternative metabolic pathways, modification of the target site, and enzymatic alteration of the antibiotic (Urban-Chmiel *et al*., 2022).

One of the most important and prevalent resistance mechanisms exhibited by Gram-negative infectious bacteria is the production of *β*-lactamases, which hydrolyse *β*-lactam rings and thereby degrade antibiotics (Tooke *et al*., 2019; Wilke *et al*., 2005; Cantòn *et al*., 2012). *β*-lactam antibiotics disrupt peptidoglycan cross-linking during cell wall synthesis, making the cell susceptible to lysis and death. Extensive and improper use of *β*-lactam antibiotics in human health and agriculture has increased the selective pressure leading to a growing number of resistant bacterial strains (Bradford, 2001; Bonnet, 2004). Successful proliferation of bacteria, with a wide multidrug resistance profile, has been further supported by Horizontal Gene Transfer (HGT), wherein mobile genetic elements like plasmids and transposons harbouring antibiotic resistance genes are exchanged between bacteria (Van Hoek *et al*., 2011). Thus, both the natural selection of mutants with high substrate specificity and the transfer of resistance genes from the surrounding metagenome lead to the evolution of *β*-lactamases with an extended-substrate spectrum (ESBLs) in the Enterobacteriaceae family (D’Andrea *et al*., 2013). ESBLs efficiently hydrolyse penicillins, as well as broad-spectrum cephalosporins (such as cefotaxime, ceftriaxone, ceftazidime) and monobactams (e.g. aztreonam), but do not effectively hydrolyse cephamycins or carbapenems and can be inhibited by some inhibitors (Cantòn *et al*., 2012; Van Hoek *et al*., 2011).

In Enterobacteriaceae, the CTX-M *β*-lactamases have largely surpassed other Ambler class A ESBL enzymes in terms of ubiquity (Cantòn *et al*., 2012). The CTX-M family consists of five main clusters: CTX-M-1, CTX-M-2, CTX-M-8, CTX-M-9 and CTX-M-25, which differ from each other by an amino acid sequence of *≥*10% (Cantòn *et al*., 2012). Out of the five, CTX-M-1 and CTX-M-9 are the most widespread clusters. The focus of the present study is CTX-M-14 (Fig. 1 a), which belongs to the CTX-M-9 subgroup and is one of the most important and hence, extensively studied enzymes of the CTX-M family (D’Andrea *et al*., 2013). CTX-M-14 *β*-lactamases can effectively hydrolyse most cephalosporins and penicillins; they hydrolyse carbapenems too, albeit with a reduced efficiency (Ishii *et al*., 2007). Essentially, Class A enzymes hydrolyse the peptide-mimicking bond in the *β*-lactam ring of the antibiotic, by stepwise acylation and deacylation. During the first part of the reaction, the active site residue Ser 70 attacks the carbonyl carbon of the *β*-lactam, cleaving the amide bond, which protonates the *β*-lactam nitrogen, forming an acyl-enzyme intermediate. In the second part of the reaction, Glu 166 then activates a catalytic water, which attacks the carbonyl carbon of the ester bond between the oxygen of Ser 70 and the *β*-lactam ring. In turn, this results in hydrolysis and release of the inactivated antibiotic and regeneration of the catalytically competent enzyme (Ambler, 1980; Drawz & Bonomo, 2010; Hata *et al*., 2006). Consequently, modification of either Ser 70 or Glu 166 impairs the catalysis and aids in trapping intermediates along the reaction coordinate pathway.

**Figure 1:**
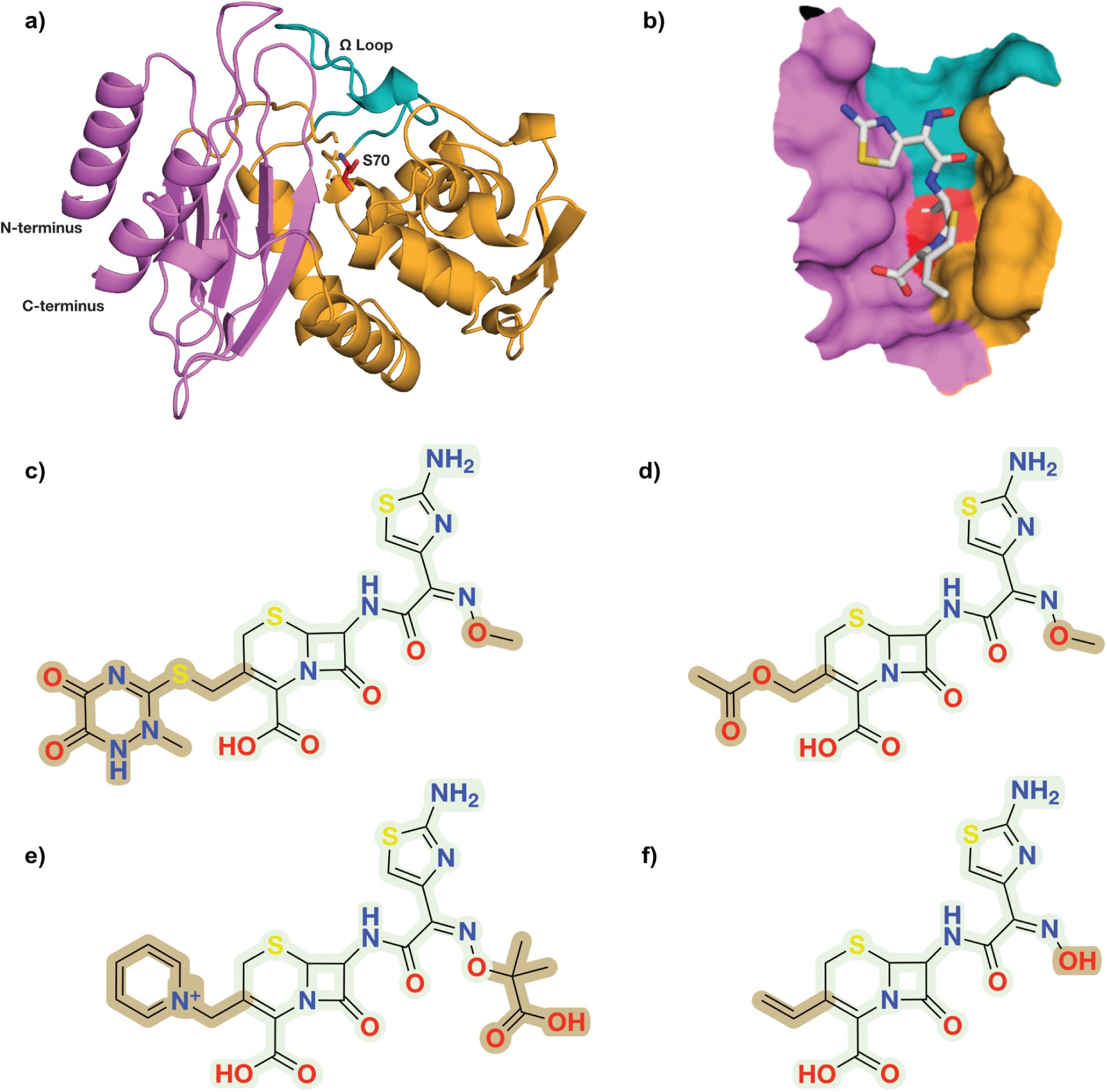
Model system, target ligand and structural homology between selective cephalosporins. a) CTX-M-14 *β*-lactamase. The catalytically important active site residue Ser 70 and the functionally relevant Ω loop (residues 160-180) are highlighted. b) Cefdinir in the active site of CTX-M-14 E166A c) Ceftriaxone d) Cefotaxime e) Ceftazidime f) Cefdinir. Differences in the side chain are highlighted.

Third-generation cephalosporins are the most commonly prescribed cephalosporins against infections caused by both Gram-positive and Gram-negative bacteria. These antibiotics consist of a six-membered dihydrothiazine ring fused to the beta-lactam ring. The C3, C4, and C7 functional groups determine their antimicrobial activity and the chemical properties (El-Shaboury *et al*., 2007). Cefdinir is a semi-synthetic, extended-spectrum oximino-cephalosporin, which has been prescribed against bacterial pneumonia, other respiratory tract infections, otitis media, and skin infections since the late 1990s (Guay, 2002). Though Cefdinir is more susceptible to hydrolysis by *β*-lactamases than larger cephalosporins such as Ceftazidime and Ceftriaxone, it remains clinically relevant due to its oral administrability.

Originally identified as Cefotaximases, CTX-Ms readily hydrolyse smaller oximino-cephalosporin derivatives, but perform poorly against bulkier varieties like Ceftazidime and Ceftriaxone. However, recent studies reveal that naturally occurring single amino acid substitutions in CTX-Ms, such as A77V, N106S, P167S/T/Q, D240G, result in a drastic increase in the cephalosporin-hydrolysing ability (Bonnet, 2004; Chen *et al*., 2005; Novais *et al*., 2008; Both *et al*., 2017; Patel *et al*., 2018). It is therefore crucial to determine the exact nature of the interaction between enzyme and substrate with clinically relevant cephalosporins, which could be helpful for future structure-based drug development.

Previously determined complexes of Cefdinir homologues like Ceftazidime and Ceftriaxone have provided invaluable insight into the ESBL active site plasticity (Brown *et al*., 2020; Patel *et al*., 2017; Patel *et al*., 2018; Lu *et al*., 2023). Recent studies have also explored the idea of Cefdinir-inhibitor combination for an improved bactericidal efficiency against *β*-lactamases (Srivastava *et al*., 2021). However, there is currently no reported crystallographic analysis of any *β*-lactamase in complex with the clinically relevant cephalosporin Cefdinir. To address this gap, we utilised serial cryo-crystallography and determined the structure of wildtype apo CTX-M-14, apo CTX-M-14 E166A, and Cefdinir bound to CTX-M-14 E166A.

## 2 Materials and method

### 2.1 Protein purification and crystallisation

The CTX-M-14 E166A gene was synthesised and cloned in a pET-24a(+) vector (BioCat GmbH, Heidelberg, Germany) with a Kanamycin selection marker. The plasmid was transformed into *E. coli* strain BL21 (DE3) and grown in LB (Luria Miller) medium supplemented with 50 µg/mL Kanamycin at 37 °C until an OD_600_ of 0.6 - 0.8 was reached. Protein expression was induced by the addition of IPTG to a final concentration of 175 µM, after which the cells were incubated at 37 °C for 4 h. The cells were harvested by centrifugation (5500 *×* g, 10 min, 4 °C) and pellets were stored at -20 °C until purification. For purification, the cell pellet was resuspended in purification buffer (20 mM MES, pH 6), followed by sonication for cell lysis. Cell debris was separated by centrifugation (20000 *×* g, 1 h, 4 °C). The cleared supernatant was dialysed overnight against a large volume of purification buffer at 4 °C using a 6-8 kDa molecular weight cut-off membrane. The CTX-M-14 protein was purified using cation exchange chromatography (5 ml HiTrap SP FF, Cytiva) and eluted using a gradient of 20 mM MES, pH 6, 0–50 mM NaCl over 5 column volumes. The protein was then concentrated to 22 mg/ml using 10 kDa centrifugal filter units (Amicon Ultra-15). For crystallisation of the activity-impaired CTX-M-14 mutant E166A, protein solution (22 mg/ml) was mixed with 45% (v/v) crystallising agent (40% (w/v) PEG 8000, 200 mM LiSO_4_, 100 mM sodium acetate, pH 4.5) and with 5% (v/v) undiluted seed stock solution to induce micro-crystallisation. This resulted in crystals with a homogeneous size distribution of ca. 11-15 µm overnight.

For soaking experiments, Cefdinir was dissolved in the stabilisation buffer (28% (w/v) PEG 8000, 140 mM LiSO_4_, 70 mM sodium acetate, 6 mM MES pH 4.5) to reach a concentration of about 50 mM. The ligand did not dissolve completely; hence, the supernatant was mixed with the crystal slurry in a (1:3) ratio with 12% (v/v) 2,3-butanediol as cryo-protectant.

### 2.2 Structure determination

About 2 *µ*l microcrystalline sample was pipetted onto a SPINE-standard mesh-loop with 400 *µ*m mesh area having 10 *µ*m openings (MiTeGen, MicroMesh). The initially established cryogenic serial data collection involved raster scanning with rotational exposure (Gati *et al*., 2014). However, we collected still diffraction data with a previously described workflow (Mehrabi *et al*., 2023) available in MXCube (Oscarsson *et al*., 2019) at the P14 beamline of PETRA-III (EMBL Unit Hamburg, Germany).

Briefly: A micro-focused 7 *×* 3 *µ*m sized (at the FWHM) X-ray beam with an energy of 12.7 keV (0.9763 Å), flux of 2 *×* 10^13^ ph/s, and an exposure time of 7.5 ms per image was used during data collection with an Eiger2 CdTe 16M detector (Dectris, Baden-Daettwil, Switzerland). During data collection, a grid with spacing matching the dimensions of the beam was drawn over the whole micro-mesh sample, giving rise to several thousand still diffraction images, which were processed using CrystFEL with the XGANDALF indexing routine (Gevorkov *et al*., 2019; White *et al*., 2012). Structures were solved by molecular replacement in PHASER using PDB entry 6GTH as a search model for CTX-M-14 (McCoy *et al*., 2007). Structure refinement was done with iterative cycles of *phenix.refine* and manual model building in COOT-v0.9 (Adams *et al*., 2004; Emsley & Cowtan, 2004; Emsley *et al*., 2010). Figures were generated with PyMOL (Schrödinger, LLC, 2015) and ChemDraw (Revvity Signals Software, Inc., Waltham, USA). Table 1 summarises data collection and refinement statistics.

**Table 1:** Data collection and refinement statistics. Values in the highest resolution shell are shown in parentheses.

| Structure (PDB-ID) | Wildtype Apo (9TKX) | E166A Apo (9TKY) | Cefdinir (9TLD) |
| --- | --- | --- | --- |
| **Data collection** |  |  |  |
| Space group | P3<sub>2</sub>21 | P3<sub>2</sub>21 | P3<sub>2</sub>21 |
| Cell dimensions |  |  |  |
| *a*, *b*, *c* (Å) | 41.82 41.81 232.36 | 41.78 41.76 232.92 | 41.82 41.82 233.09 |
| $\alpha$, $\beta$, $\gamma$ (°) | 89.97 89.99 119.98 | 89.97 89.99 120.00 | 89.97 89.98 119.99 |
| Resolution range (Å) | 116.28 - 1.63<br>(1.66 - 1.60) | 116.38 - 1.63<br>(1.66 - 1.60) | 116.28 - 1.63<br>(1.66 - 1.60) |
| Diffraction patterns | 58030 | 81966 | 30491 |
| Total reflections | 74240034<br>(5006769) | 85504320<br>(5774126) | 43172014<br>(2903342) |
| Unique reflections | 32733 | 32733 | 32733 |
| Redundancy | 2268(1572.5) | 2612.2 (1813.5) | 1318.9 (911.9) |
| Completeness (%) | 100 (100) | 100 (100) | 100 (100) |
| Mean I/$\sigma$ (I) | 17.57 (8.72) | 21.94 (12.07) | 13.47 (11.03) |
| Wilson B-factor (Å<sup>2</sup>) | 17.55 | 15.43 | 18.07 |
| *R*<sub>split</sub> | 0.680 (1.431) | 0.677 (1.123) | 1.181 (1.061) |
| CC1/2 | 0.993(0.974) | 0.990 (0.986) | 0.981 (0.973) |
| CC* | 0.998(0.993) | 0.998 (0.997) | 0.996 (0.993) |
| **Refinement** |  |  |  |
| Reflections used in refinement | 32597 (2491) | 32588 (2487) | 32591 (2766) |
| *R*<sub>work</sub> | 0.2291 (0.2737) | 0.2310 (0.2934) | 0.2103 (0.1811) |
| *R*<sub>free</sub> | 0.2537 (0.3125) | 0.2555 (0.3584) | 0.2464 (0.2259) |
| Reflections used for *R*<sub>free</sub> | 1612 (134) | 1616 (134) | 1543 (140) |
| Number of non-hydrogen atoms | 2352 | 2351 | 2293 |
| macromolecules | 2119 | 2051 | 1987 |
| ligands | 5 | 5 | 26 |
| solvent | 228 | 295 | 280 |
| Average B-factor (Å<sup>2</sup>) | 21.37 | 21.86 | 19.02 |
| macromolecules | 20.52 | 20.79 | 17.87 |
| ligand | 28.83 | 21.41 | 20.91 |
| solvent | 29.09 | 29.29 | 27.02 |
| **RMS deviations** |  |  |  |
| Bond lengths (Å) | 0.005 | 0.010 | 0.007 |
| Bond angles (°) | 0.869 | 1.187 | 0.953 |

## 3 Results and Discussion

### 3.1 Cryo-SSX

Serial synchrotron crystallography at cryo-conditions (cryo-SSX) is a useful alternative to conventional single-crystal cryo-crystallography when large crystals are difficult to produce, but micro-crystals are available. When solubility issues with the ligand result in problematic soaking times for larger crystals, microcrystals can be a viable alternative, thanks to their favourable surface-to-volume ratio. Another significant advantage is the reduction in radiation damage that can be achieved through serial data collection, whereby the dose is distributed across thousands of crystals.

Unlike SSX at room temperature, cryo-SSX offers the advantage of standardised sample handling: samples can be prepared prior to the beamtime, loaded onto commercially available holders and shipped. This standardisation enables access to high-throughput data collection workflows, such as those found at many synchrotron beamlines.

On the other hand, cryo-SSX poses sample-related challenges: too high a crystal density may result in overlapping diffraction patterns, while conversely, too low a crystal density may result in an inadequate orientational multiplicity. Such a lack of orientational multiplicity among the indexed microcrystals might cause inferior crystallographic data quality metrics (e.g. R-factors) during model refinement, than the resolution of the dataset would imply for canonical rotation data. Unfortunately, a clear metric indicating said orientational bias is currently missing. While further data analysis tools will be required in the future to ascertain the cause, we have attempted to circumvent these problems by careful sample preparation and sampling a larger number of diffraction patterns than typically required for room-temperature SSX (Moreno-Chicano *et al*., 2019; Gorel *et al*., 2021; Mehrabi *et al*., 2021).

Here we have determined the structure of the ESBL CTX-M-14, its activity-impaired mutant E166A, and of CTX-M-14 E166A in complex with the third-generation extended-spectrum cephalosporin Cefdinir (Fig. 1, 2), via cryo-SSX. During data collection, we obtained 105,490 diffraction images for the wild-type apo structure, of which 58,030 were indexable. Similarly, out of the 157,342 diffraction images of the E166A apo structure, 81,966 were indexable. Along those lines, we obtained 61,422 diffraction patterns for the Cefdinir-bound complex, of which 30,491 could be successfully indexed. All structures were solved by molecular replacement with one monomer in the asymmetric unit and were refined to comparable and reasonable data quality parameters (Table 1).

### 3.2 Influence of the E166A mutation on the apo structure of CTX-M-14

The activity-impaired CTX-M-14 mutant E166A crystallised with one monomer per asymmetric unit. The protein adopts the canonical, previously observed two-domain structure (Lee *et al*., 2025). Superposition with the wildtype protein demonstrates a root mean square deviation (RMSD) of 0.15, thereby substantiating the observation of the overall high structural agreement between the mutant and the wildtype enzyme.

At the site of the mutation, the backbone structures can be superimposed with minimal differences. One significant difference between the WT and E166A mutant is the position of the catalytic water molecule essential for the successful completion of ligand hydrolysis. In case of the E166A mutant, the catalytic water, otherwise present in the WT near the active Ser 70, is displaced *≈*1.5 Åaway due to a lack of coordination with Glu 166. This, in turn, prevents the de-acylation step and traps the acyl-enzyme intermediate. The E166A mutation seems to induce a slight backbone shift in residues Thr 165 – Leu 169, especially in Pro 167 (0.4 Å). Nevertheless, this shift does not prevent Cefdinir from being placed in its current position, supporting the hypothesis that the reduction in catalytic activity can be assigned to functional rather than structural differences between the proteins.

### 3.3 CTX-M-14 E166A Cefdinir interaction

After molecular replacement of the E166A-Cefdinir complex, we determined strong difference electron density in the active site. The electron density encompasses the entire ligand and is clearly connected to the active site Ser 70, indicating a hydrolysed, covalently bound Cefdinir, resembling the acyl-enzyme intermediate.

Cefdinir, as characteristic of cephalosporins, contains a core where a four-membered *β*-lactam ring is fused to a six-membered dihydrothiazine ring. The C3 position of the core is extended via a vinyl group, while the C4-carbon of the core is attached to a carboxyl group. Additionally, the C7-carbon of the core is bound to an aminothiazolyl-oxime-acetamido sidechain (Fig. 1 c).

Similar to the interactions observed in previously determined CTX-M-14 substrate complexes, Cefdinir also interacts with the typical active site residues of CTX-M-14. Canonically, ligands bound to CTX-M-14 active sites are stabilised by a hydrogen bond network with the surrounding Ser 70, Asn 104, Ser 130, Asn 132, Pro 167, Asn 170, Thr 235, Ser 237, and Asp 239 residues, which is also reflected in the Cefdinir complex (Fig 2). Usually, hydrogen bonds between the carbonyl oxygen of the *β*-lactam and the main chain nitrogens of Ser 70 and Ser 237 stabilise the developing negative charge on the tetrahedral intermediate during acylation (Matagne *et al*., 1998). This pattern is also observed in the Cefdinir complex, where 2.8 Å and 2.9 Å hydrogen bonds are maintained between the carbonyl oxygen and main chain nitrogens of Ser 70 and Ser 237, respectively. The Ω loop encloses the active site in Class-A *β*-lactamases (Fetrow, 1995; Ibuka *et al*., 1999; Poirel *et al*., 2001). Important interactions necessary to preserve the structural integrity of the Ω loop (Ibuka *et al*., 2003; Chen *et al*., 2005) are also conserved in the Cefdinir complex. For instance, salt bridges between Arg 164 and Asp 179, Arg 161 and Asp 163, Asp 176 and Arg 178 are present. Additionally, hydrogen bonds between Arg 164 and Thr 171, Ala 166 and Asn 136 (contrary to the hydrogen bond between Glu 166 and Asn 170 in the wildtype enzyme) are also observed. Thus, the Ω loop is well preserved in the E166A mutant and shows no significant conformational variation upon ligand binding. Additionally, the conformation of loops 103-106 and 213-220 has been reported to be important for efficient binding and hydrolysis of cephalosporins (Patel *et al*., 2018; Lu *et al*., 2023). The crystal structure shows that these loops are well preserved in the CTX-M-14 E166A-Cefdinir complex.

**Figure 2:**
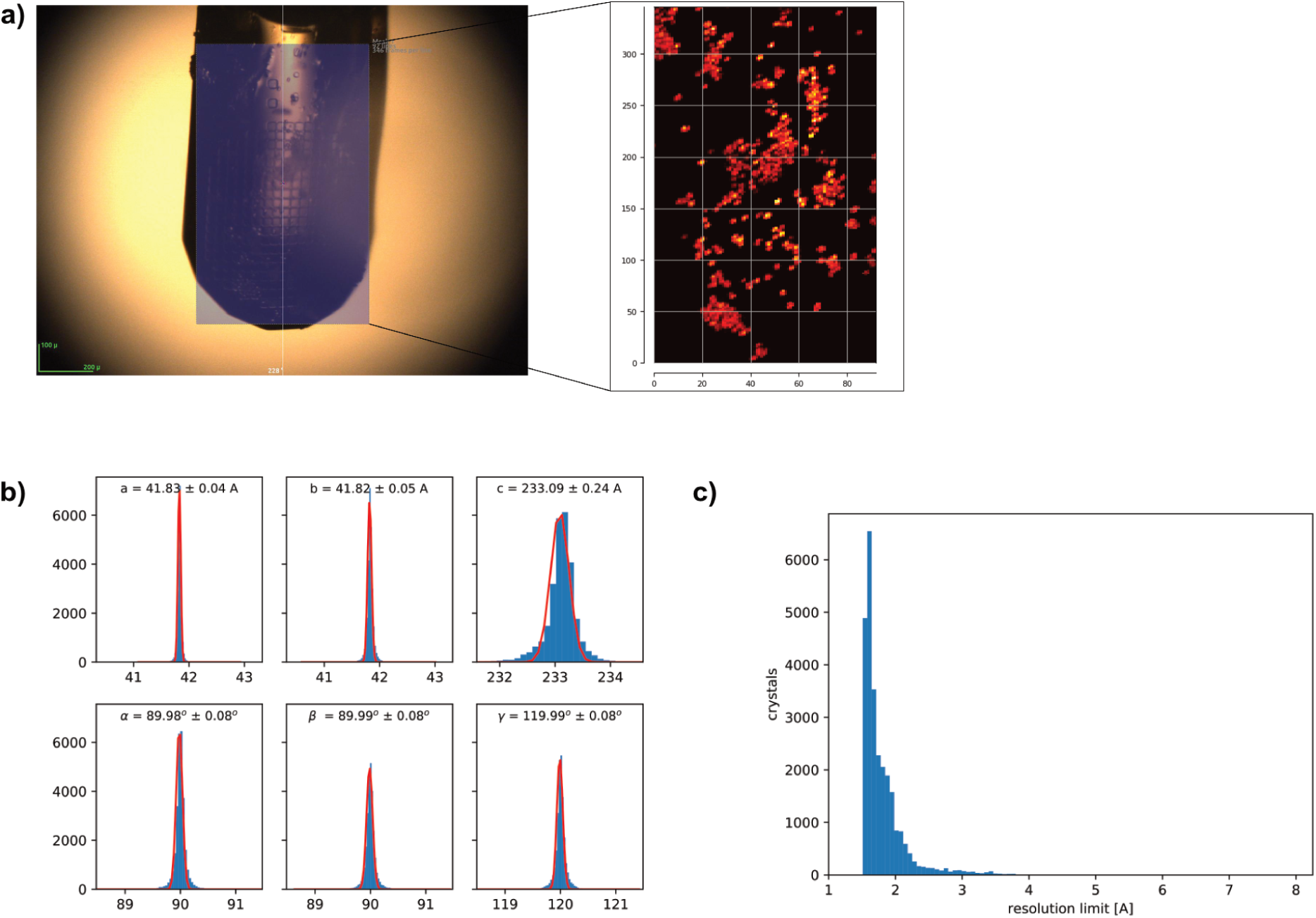
Selective example of typical serial cryo data collection and processing output. a) Left panel shows the drawn raster scanning grid onto the SPINE-standard mesh loop with microcrystals. The right panel shows the corresponding heat map of crystal hits. b) Unit cell distribution of microcrystals. c) Resolution distribution of identified microcrystals.

### 3.4 Comparison to other Cephalosporins

Chemically, the differences between broad-spectrum third-generation cephalosporins present in the PDB, such as Cefdinir, Ceftriaxone, Cefotaxime, and Ceftazidime, are located in the C3 and C7 sidechains (highlighted in Fig. 1, c-f). Cefdinir has a vinyl group (C3) and an aminothiazolyloxime-acetamido sidechain (C7) (Fig. 1 c). Ceftriaxone has the thiotrizinedione (C3) and an aminothiazole-methoxyimino group (C7) (Fig. 1 d), Cefotaxime has an acetoxymethyl group (C3) and an aminothiazole-methoxyimino group (C7) (Fig. 1 e) and finally, Ceftazidime has a pyridinium group (C3) and an aminothiazole-methoxyimino group with a carboxypropyl tail (C7) (Fig. 1 f) To understand how Cefdinir compares to other cephalosporin derivatives, we compared our structure to CTX-M-14 complexes with Cefotaxime (Soeung *et al*., 2020) (PDB-ID: 7K2W, E166A/K234R mutant) and Ceftazidime (Patel *et al*., 2017) (PDB-ID: 5U53, E166A mutant). Previously determined Ceftriaxone is bound to BlaC, and thus is excluded from the present comparison.

To understand global structural differences between the different ligand complexes, we calculated the C*_α_*-RMSD between wildtype apo, apo E166A, E166A/K234R:Cefotaxime, E166A:Ceftazidime and E166A:Cefdinir (Fig 4, a). The categorical heatmap shows C*_α_*-RMSD values with a maximum *≈* 0.3 Å, which indicates no significant backbone changes between the complexes and is reflected in a good superposition of the structures (Fig. 4 b). As the heatmap indicates, the structures overlay quite well. The structurally and functionally important Ω loop does not show pronounced differences when aligned. The only striking difference between the five is the loop enclosing residues 252-257, which are not part of the canonical catalytically important residues. Interestingly, our two SSX apo structures and the acyl-enzyme complex align very well in this region, whereas the Cefotaxime and Ceftazidime datasets differ from each other as well as our structures. Cryo trapping or differences in crystallisation conditions could have introduced the variation.

The CTX-M-14 Cefotaxime and Ceftazidime complexes also exhibit interactions similar to the Cefdinir complex, Fig. 3 c. Superposition of the ligands in the active site of CTX-M-14 (Fig. 4, c) indicates a partially flipped binding pose of Ceftazidime. This different ligand orientation may be caused by the slightly bulkier C7 sidechain in Ceftazidime. It’s worth noting that the C3 sidechains of Cefotaxime (acetoxymethyl) and Ceftazidime (pyridinium) have likely been eliminated during the acylation and formation of the acyl-enzyme intermediate, as observed in other serine *β*-lactamase structures in complex with cephalosporin antibiotics (Hamilton-Miller, 1994; Patel *et al*., 2017; Tooke *et al*., 2021). However, there was no elimination in the case of Cefdinir, even though an acyl-enzyme intermediate was formed. Likely, the presence of a vinyl sidechain rather than a good leaving group at C3 prevents fragmentation.

**Figure 3:**
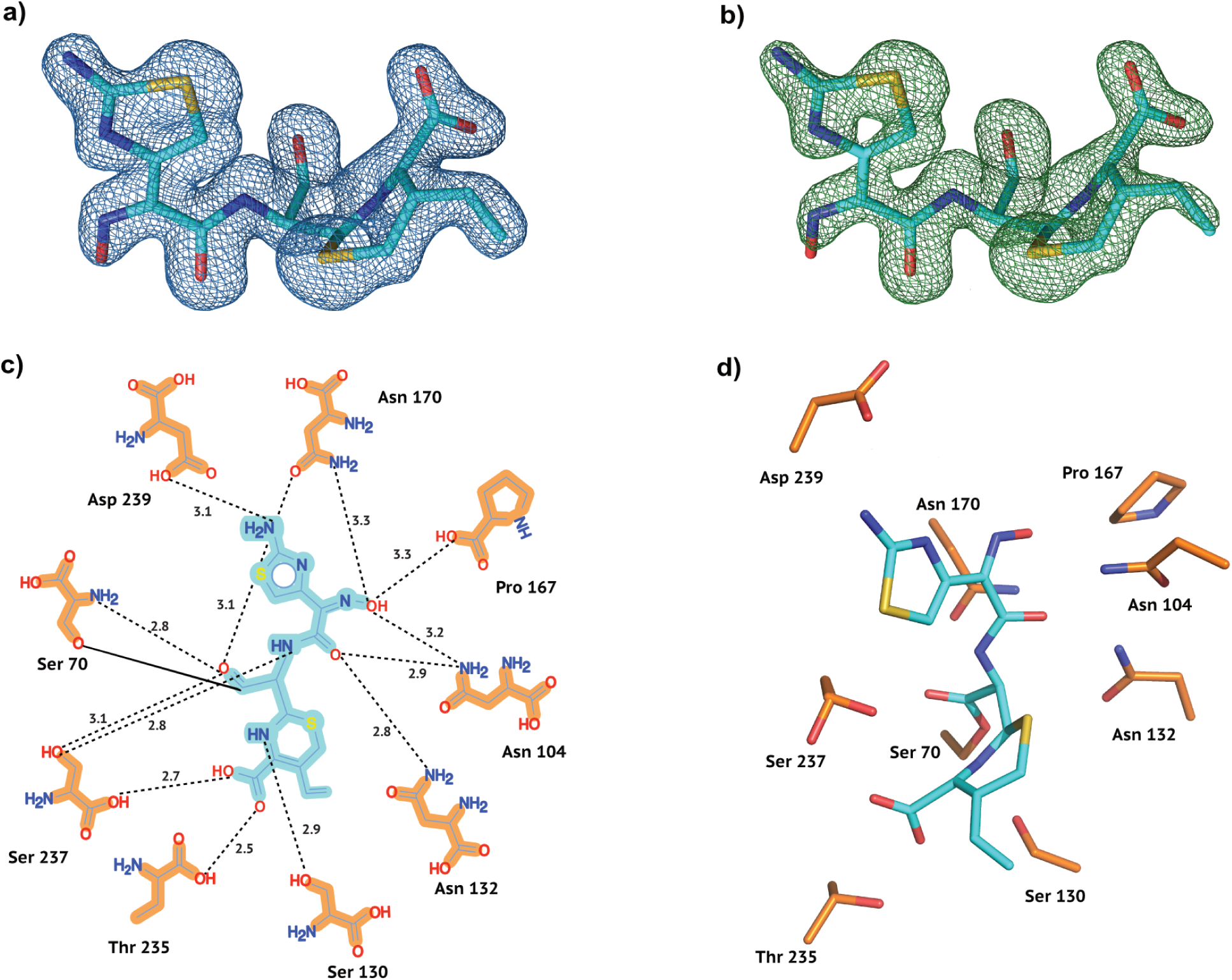
Electron density and conformation of Cefdinir at cryo temperature. a) 2F*_o_*-F*_c_* map shown at an RMSD of 1.0. b) POLDER omit map shown at an RMSD of 3.0. c) 2D projection of Cefdinir and the contact residues showing interatomic distances. d) Cefdinir conformation in the CTX-M-14 active site. Hydrogen bonds are represented as black dotted lines with their distance given in Å.

**Figure 4:**
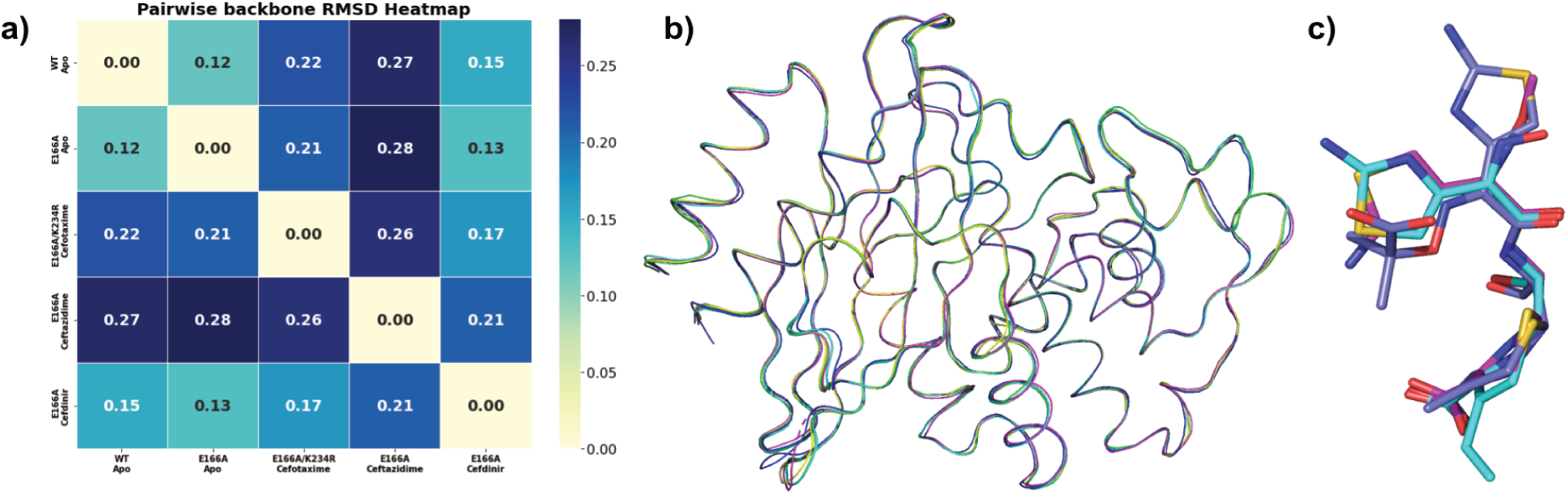
Structural comparison of homologues. a) Pairwise C*α* categorical heatmap. b) Ribbon representation of superposed wildtype apo (yellow), E166A apo (green) Cefotaxime (magenta, PDB ID: 7K2W), Ceftazidime (blue, PDB ID: 5U53) and Cefdinir (cyan, PDB ID: ) c) Superposed Cefotaxime (magenta, PDB ID: 7K2W), Ceftazidime (blue, PDB ID: 5U53) and Cefdinir (cyan, PDB ID: 9TLD)

### 3.5 Conformational differences induced by Cefdinir binding

A difference can be observed between the three (Fig. 5 a) acyl-enzyme intermediate complexes (Fig 5 a, b, c), around Ser 130, a residue contributing to both substrate binding and catalysis of cephalosporins (Lu *et al*., 2023). In case of E166A:Ceftazidime and E166A:Cefdinir complexes, Ser 130 forms hydrogen bonds with Lys 73, Lys 234 (which in turn interacts with a water molecule and Thr 235), and the dihydrothiazine ring nitrogen. In the E166A/K234R:Cefotaxime complex, as a result of the K234R mutation, Ser 130 is reoriented by almost 90° to form a hydrogen bond with Arg 234, losing the contact with Lys 73, but in turn, hydrogen bonds with the carboxylate oxygen of the ligand.

**Figure 5:**
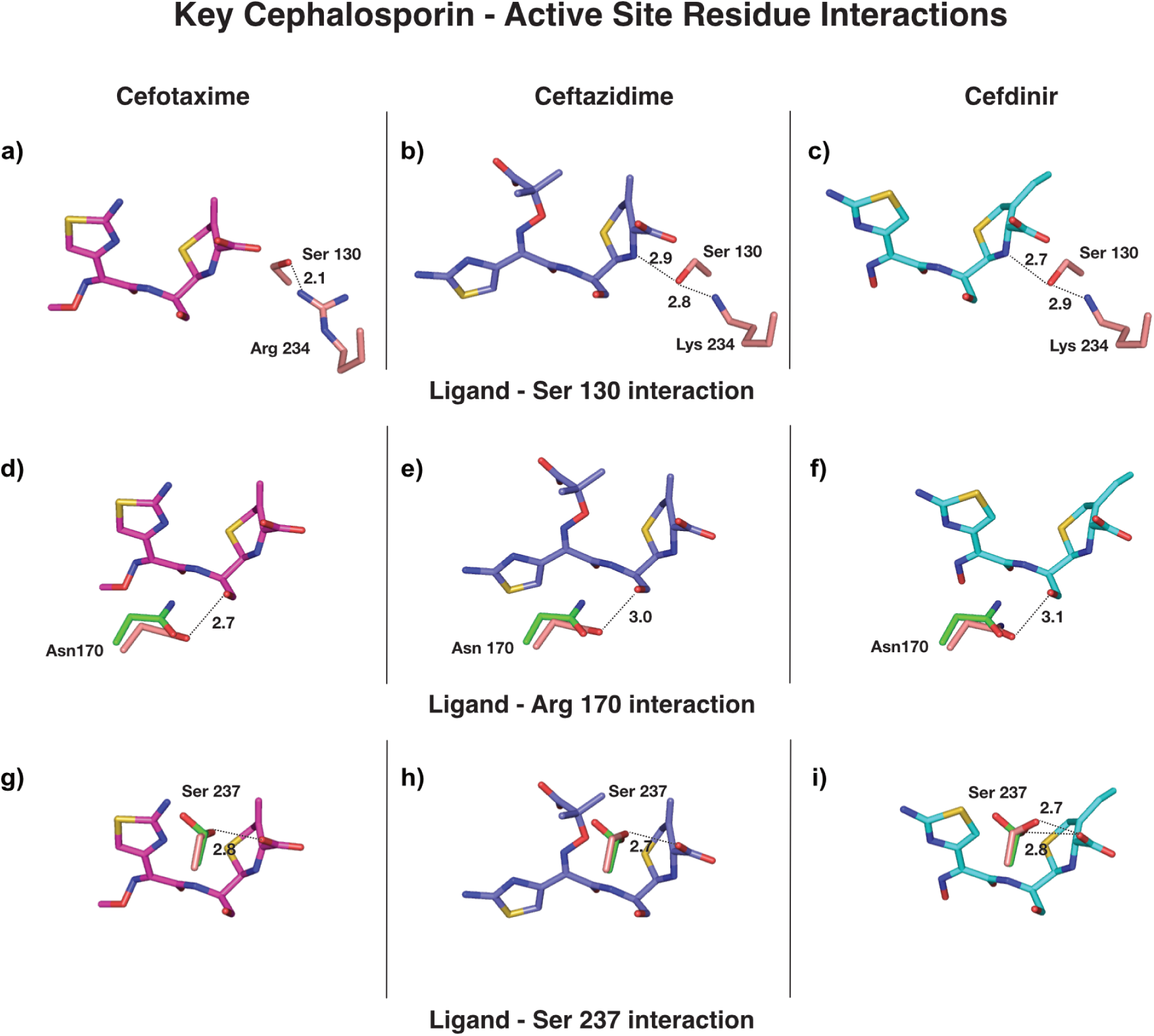
Comparison of key ligand-residue interactions in Cefotaxime, Ceftazidime and Cefdinir. a) Residues S130 and K234 interacting with Cefotaxime b) Residues S130 and K234 interacting with Ceftazidime c) Residues S130 and K234 interacting with Cefdinir d) Residue N170 interacting with Cefotaxime e) Residue N170 interacting with Ceftazidime f) Residue N170 interacting with Cefdinir g) Residue S237 interacting with Cefotaxime h) Residue S237 interacting with Ceftazidime i) Residue S237 interacting with Cefdinir

Fig. 5 d,e,f show Asn 170 from each complex (salmon) superposed with Asn 170 from the apo E166A (green) structure. It was hypothesised that in CTX-Ms, the hydrogen bond between Ω loop residue Asn 170(O) and Asp 240(N) breaks to widen the active site to accommodate cephalosporins like Cefotaxime into the binding pocket (Adamski *et al*., 2015; Delmas *et al*., 2010). Thus, this hydrogen bond remains intact in the wild-type apo CTX-M-14 but is disrupted in substrate-bound structures.

Similar to the Cefotaxime and Ceftazidime complexes, this bond is missing in the Cefdinir complex, indicating that Cefdinir leads to a widening of the active site, too. As a consequence of this, the Asn 170 sidechain orientation is slightly altered in cephalosporin-bound structures compared to the apo WT and CTX-M-14 E166A enzymes. In all cephalosporin-bound structures, Asn 170 is positioned to interact with the carboxyl oxygen of the *β*-lactam.

Conservation of residues Ser 237 and Arg 274 throughout the CTX-M family and corresponding published studies have revealed that the two residues work in synchrony to stabilise bound cephalosporins (Adamski *et al*., 2015; Pérez-Llarena *et al*., 2008; Gazouli *et al*., 1998*a*; Delmas *et al*., 2010; Gazouli *et al*., 1998*b*). The sidechain of Ser 237 interacts with the C4 carboxyl of the ligand in all three cases (Fig. 5 g, h, i). The figures also show apo E166A Ser 237 (green) superposed to the Ser 237 of the ligand-bound enzyme (salmon). The apo Ser 237 assumes two conformations; one is clearly directed about 150° away from the orientation resembling Ser 237 of the ligand-bound enzyme. Unlike in the Cefotaxime and Ceftazidime complexes, in the E166A-Cefdinir complex, we observe two orientations of the Ser 237 sidechain. This implies an intrinsic mobility of Ser 237, potentially important for substrate binding and stabilisation.

To conclude, our study reveals the acyl-enzyme complex between cephalosporin Cefdinir and CTX-M-14 E166A obtained via cryo-SSX. Comparison with structural homologues confirms a comparable binding mode. Moreover, considering that currently only about 5% of all cryo protein structures in the PDB are obtained via SSX, this study emphasises that cryo-SSX is a valid alternative to canonical single-crystal data-collection, and can be exploited for the reliable characterisation of protein-ligand interactions.

## Acknowledgements

All data were collected at beamline P14 operated by EMBL Hamburg at the PETRA-III storage ring (DESY, Hamburg, Germany). We would like to thank our colleagues A.R. Pearson and P. Mehrabi for their continuous support and helpful discussions. E.C.S. designed the experiments; G.G. and E.C.S. performed the data collection with support from D.v.S.; A.P. and K.B. prepared the protein and the protein crystals; G.G., D.v.S., and E.C.S. processed and analysed the diffraction data. G.G. refined the structures. G.G. and E.C.S. wrote the manuscript; All authors discussed and corrected the manuscript.

## Data availability

All coordinate and structure factor files have been released in the protein data bank under PDB-IDs: 9TKX, 9TKY and 9TLD

